# Mechanism of Alkaline Gating in a Pentameric Ion Channel

**DOI:** 10.64898/2026.01.07.698087

**Authors:** Emelia Karlsson, Ida Ygland, Anton Jansen, Jeffrey Plumley, Erik Lindahl, Rebecca J. Howard, Berk Hess

**Affiliations:** SciLifeLab, Department of Biochemistry & Biophysics, Stockholm University, PO Box 1031, SE-171 21 Solna, Sweden; SciLifeLab & Swedish e-Science Research Center, Department of Applied Physics, KTH Royal Institute of Technology, PO Box 1031, SE-171 21 Solna, Sweden; University of Oregon, Eugene, OR 97403, USA; Department of Physics, Chemistry and Biology, Linköping University, SE-581 83 Linköping, Sweden

**Keywords:** sTeLIC, ligand-gated ion channel, electrophysiology, constant-pH molecular dynamics simulation, titratable residues, pH activation, proton sensor

## Abstract

Pentameric ligand-gated ion channels (pLGICs) are key mediators of electrochemical signal transduction in various organisms. Like many proteins involved in cellular signaling, they are modulated by a variety of environmental factors including pH and small molecules. However, the molecular mechanisms underlying pLGIC activation and modulation remain unclear. A promising model system in this family is the bacterial ion channel sTeLIC, which can be activated by alkaline pH, and for which we recently determined structures in multiple functional states. However, protonation changes and other pH-driven dynamics cannot be directly observed in these structures. Here, we used constant-pH molecular dynamics simulations and oocyte-electrophysiology recordings from engineered mutants to develop a comprehensive mechanistic model for pH sensing. Interestingly, critical residues include two Glu residues (E106, E160) located in the extracellular-vestibule and domain-interface regions of each subunit, where they mediate differential electrostatic interactions in closed versus open states. This work demonstrates the applicability of constant-pH methods to model dynamic processes in a multimeric membrane-embedded protein, and offers a detailed mechanism for pH sensing, likely extensible to human drug targets.

## Introduction

Pentameric ligand-gated ion channels (pLGICs) are key mediators of electrochemical signal transduction in most kingdoms of life [1]. In the mammalian nervous system, pLGICs mediate passive ion transport in response to specific neurotransmitters [2], and are targets in the treatment of neurological disorders, such as epilepsy [3], Parkinson’s, and Alzheimer’s disease [4]. The discovery of prokaryotic pLGICs has further advanced our comprehension of the structure and function of these proteins, owing to their comparable topology and pharmacology [5]. Among other things, prokaryotic as well as eukaryotic pLGICs are sensitive to environmental factors including small molecules and pH, offering model systems for structure-function studies of gating and modulation across the family [6].

Despite substantial variation in sequence [5], many structural motifs and molecular architectures are conserved among prokaryotic and eukaryotic pLGICs. The minimally preserved topology consists of five subunits arranged in a pseudosymmetric manner around an ion-conducting pore. Each subunit is composed of an extracellular domain (ECD) with ten ω-strands (ω1 to ω10), and a transmembrane domain (TMD) with four ε-helices (M1 to M4) [7] (Fig. 1A). Classical neurotransmitter activation occurs via a so-called orthosteric site capped by the ω9-ω10 strands (loop C) at the ECD subunit interfaces, with additional allosteric sites of activation or modulation distributed throughout the protein (Fig. 1B). However, the structural basis for sensing environmental factors such as pH remains unclear.

**Figure 1:**
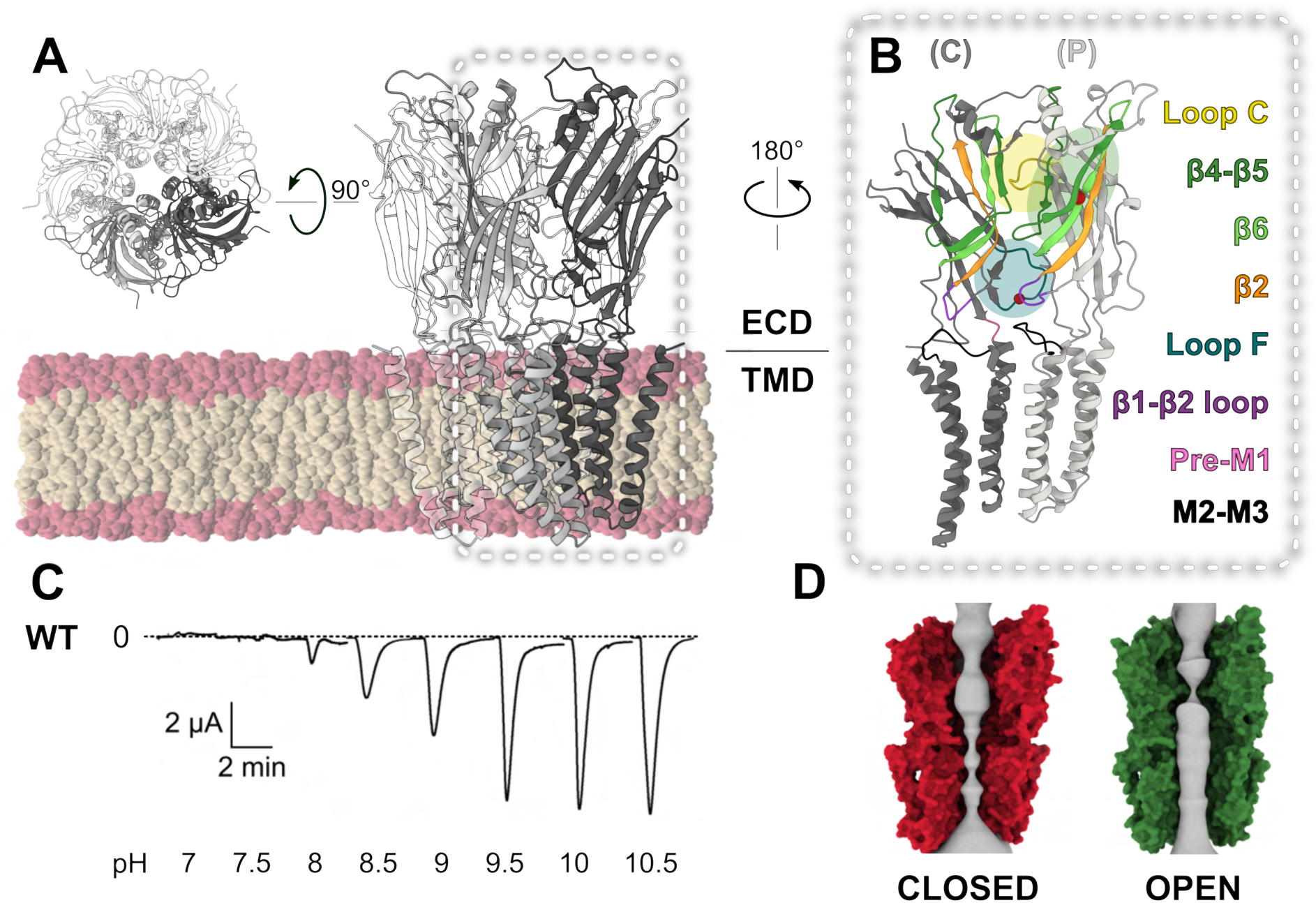
Structural and functional overview of sTeLIC. **(A)** Views from the membrane plane (left) and extracellular side (right) of a previously published closed cryo-EM structure of sTeLIC (PDB ID 9EX6 [16], gray ribbons), here embedded in a POPC lipid membrane (colored spheres). **(B)** Zoom view of the dashed region in panel A, showing the distal two sTeLIC subunits forming the principal (P) and complementary (C) faces of a single subunit interface. Shaded circles highlight the orthosteric site (yellow), ECD vestibule (green), and ECD-TMD interface (teal). Ribbon coloring in these regions indicates the ω1–ω2 loop (purple), ω2 strand (orange), ω4–ω5 loop (!-loop, dark green), ω6 strand (light green), ω8–ω9 loop (loop F, teal), and ω9–ω10 loop (loop C, yellow) in the ECD, as well as the pre-M1 (pink) and M2–M3 (orange) regions of the TMD. **(C)** Representative oocyte electrophysiology trace for WT sTeLIC, showing progressive activation by 30-s applications of increasing-pH buffers. The dashed line represents 0 µA current. **(D)** Pore surfaces (solid gray) calculated using CHAP [19] depict the water-accessible linear pathway through the ECD and the TMD in closed (PDB ID 9EX6 [16], left, red) and open structures (PDB ID 9EWL [16], right, green). For clarity, protein surfaces are shown for only the distal three subunits.

Akin to several eukaryotic receptors [8–12], some prokaryotic pLGICs are sensitive to pH. Whereas GLIC and DeCLIC are more open below neutral pH [13, 14], the channel sTeLIC, from an endosymbiont of the tube worm *Tevnia jerichonana*, activates under relatively alkaline conditions [15] (Fig. 1C). Structures of sTeLIC have been reported in a closed state at neutral pH, and an open state at elevated pH and in the presence of positive modulators [15, 16] (Fig. 1D). Notably, the extracellular vestibule mediates modulation by amphiphilic compounds [15, 16] a feature that appears to be preserved in mammalian serotonin type-3 receptors (5-HT_3A_Rs) [17]. Protonation states are not directly observable in available structures or classical molecular dynamics (MD) simulations, making it challenging to characterize mechanisms of pH gating in channels such as sTeLIC in molecular detail. However, recent advances in scalable constant-pH MD methods offer new opportunities to simulate complex protein behavior with continuous changes in atomic protonation states [18].

In this study, we employed constant-pH MD simulations and oocyte electrophysiology to investigate pH gating in sTeLIC. Recent structures of sTeLIC in both closed and open states [16] enabled quantification of protonation fractions and hydrogen-bond occupancy of titratable residues across various states and pH conditions. Putative pH sensors were then tested by oocyte electrophysiology, enabling us to identify key residues at the ECD-TMD interface and in the extracellular vestibule, and substantiating a multi-site mechanism for channel opening and ion permeation.

## Results

### State-Dependent Protonation Changes at Prospective pH-Sensing Residues

To explore pH gating in the bacterial pLGIC sTeLIC, we first performed constant-pH MD simulations of recent experimental structures in closed (PDB ID 9EX6 [16]) and open (PDB ID 9EWL [16]) states. To optimize for replicable pH-dependent differences, we ran three replicate 2-µs simulations of each system under both resting (pH 6.5) and activating (pH 10) conditions. Protonation states were allowed to vary for the side chains of all Asp, Glu, His, Tyr, Arg, and Lys residues, for which reference solution-phase p*K*_a_ values fall between 3.6 and 13.8 [20, 21], resulting in a total of 395 titratable residues in each system (79 per subunit). Throughout all simulations, the root-mean-square deviation (RMSD) of C_ε_ atoms in the protein was *<* 3 A° (Fig. S1), and the pores of closed and open structures were consistently dehydrated and hydrated respectively, indicating generally stable behavior (Fig. S2).

To identify amino-acid residues potentially involved in pH gating, we quantified the protonation fraction of each titratable side chain in sTeLIC in each simulated system (Fig. 2). We expected that residues involved in pH gating would exhibit state-as well as pH-dependent differences in protonation and/or electrostatic interactions, as previously observed for the related acid-gated pLGIC GLIC [22]. Given that sTeLIC activates upon alkalinization of the extracellular medium [15], we expected that titration of residues for which a lower likelihood of protonation is associated with the open state should facilitate activation. Specifically, we expected to find differences in protonation under relatively alkaline versus acidic pH at basic residues such as Arg, His, Lys or Tyr, whose side-chain p*K*_a_’s in solution are above 6 (Fig. 2A).

**Figure 2:**
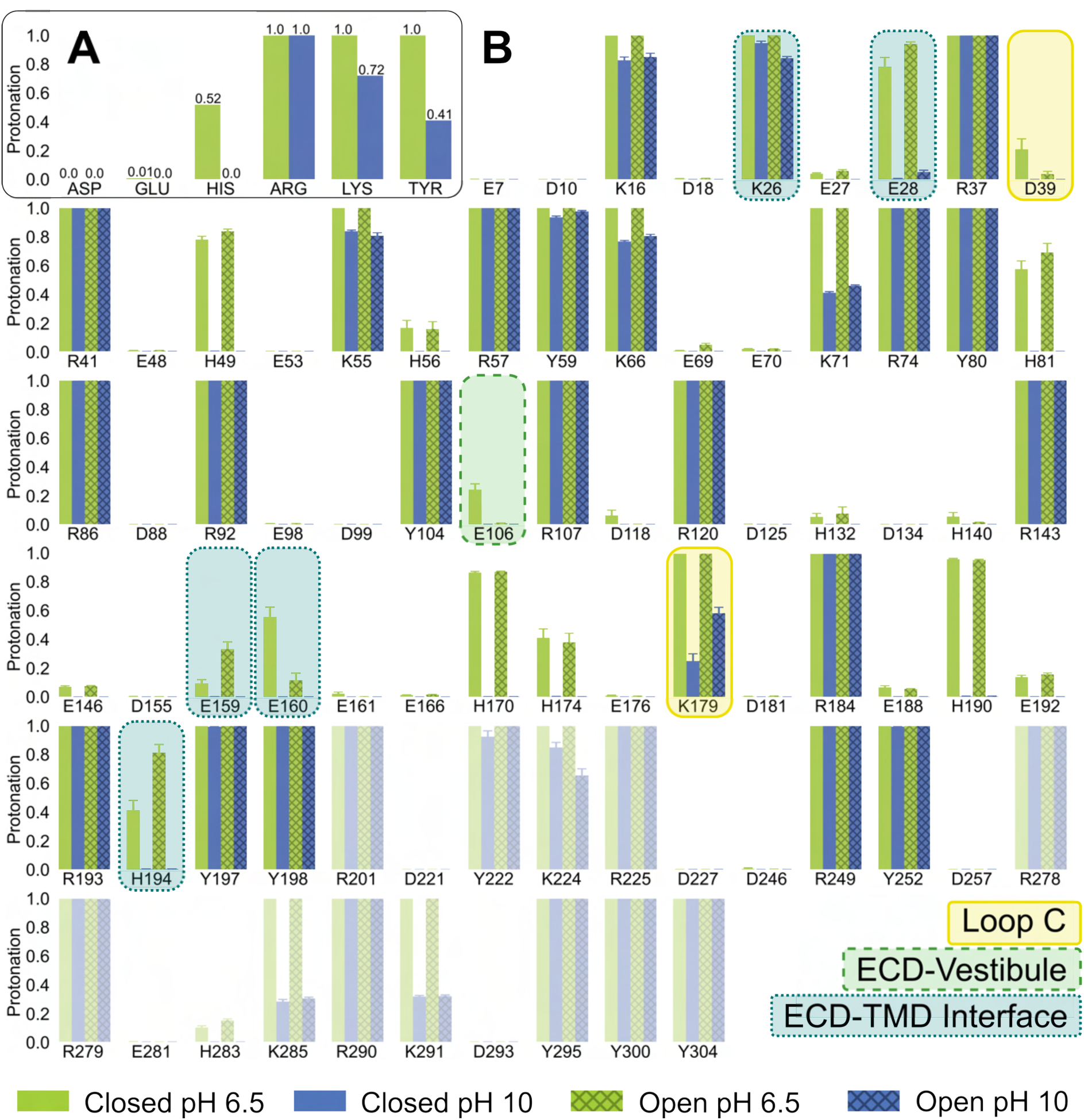
State-dependent protonation changes at prospective pH-sensing glutamates. **(A)** Protonation fractions for Asp (p*K*_a_ 3.65), Glu (p*K*_a_ 4.25), His (p*K*_a_ 6.52), Arg (p*K*_a_ 13.80), Lys (p*K*_a_ 10.40), and Tyr (p*K*_a_ 9.84) residues at resting and activating pH, based on calculations from potentiometry values for pentapeptides in solution and the Henderson-Hasselbalch equation [20, 21]. **(B)** Mean protonation fractions for each Asp, Gly, His, Arg, Lys, and Tyr residue in a single sTeLIC subunit in the closed (solid) and open (crossed) state, simulated at pH 6.5 (lime) and pH 10 (blue). Error bars indicate standard error of the mean. Residues at which mean protonation exhibits a statistically significant difference of at least 10 percentage points between open and closed simulations at the same pH (see Table S1) are colored by their location in loop C (yellow), the ECD vestibule (green), or the ECD-TMD interface (teal). Residues facing the membrane or cytosol are semi-transparent.

Notably, all 17 Arg and 10 Tyr residues in each chain were evidently protonated throughout simulations at pH 10 as well as 6.5 (Fig. 2B). Instead, most state-dependent differences in protonation appeared at acidic Asp or Glu residues, for which the protein environment evidently increased the local p*K*_a_ to enable at least partial neutralization at pH 6.5. In particular, at least 10 % less protonation was observed in the open versus closed states at equivalent pH for K26 and E160 at the ECD-TMD interface, E106 in the vestibular site, and D39 in the orthosteric site (Fig. 2B, Table S1). Accordingly, these three regions were targeted for functional characterization, as detailed in subsequent sections. Four residues (E28, E159, K179, H194) exhibited inverse state dependence, with greater protonation in open versus closed systems; these were deemed unlikely to drive alkaline gating, but were also targeted for further analysis as potentially informative controls. Six basic residues (H49, K66, K71, H170, H174, and H190) exhibited partial deprotonation at pH 10, but without clear dependence on the functional state. Three more Lys residues (K224, K285, and K291) partly deprotonated in at least one state at pH 10, but were excluded as likely mechanistic sensors of extracellular pH, due to their location at the intracellular end of the TMD. The remaining 14 Asp, 14 Glu, and 5 His residues in each chain were consistently deprotonated, while the remaining 3 Lys residues were consistently protonated, throughout simulations at pH 6.5 and 10.

Interestingly, protonation fractions observed in our constant-pH simulations were only partially consistent with predictions based on the static starting structures. Using the tool PropKa [23, 24], residues K26 and E160 at the ECD-TMD interface were already predicted to be less protonated in open versus closed starting structures, indicating this differential preference was strongly dependent on the initial configurations (Table S2). Conversely, protonation fractions at E106 and D39 were predicted to change by less than 5 % between starting structures, indicating that MD was able to capture dynamic electrostatic changes not evident in static analysis. Static-structure analysis predicted at least a 15 % decrease in protonation upon channel opening at pH 6.5 for E28, H194 and Y197 at the ECD-TMD interface, as well as H174 in the orthosteric site; these residues exhibited negligible or inverse pH sensitivity in our simulations (Fig. 2B, Table S1), suggesting their state dependence might be unstable. These residues were targeted for comparative analysis, as detailed below. Several other basic residues (H49, H81, H132, K179, K224) were predicted to protonate more in open versus closed structures, and thus appeared unlikely to drive alkaline gating. H283, Y300, and Y304 were predicted to shift in protonation upon opening, but were excluded as mechanistic sensors due to their intracellular orientation.

### Deprotonation at E160 Promotes Activation and Na^+^ Accumulation

The ECD-TMD interface undergoes notable conformational changes upon sTeLIC opening, especially around loop F on the complementary side of each subunit interface (Fig. 3A, top). In the most dramatic state-dependent difference between closed and open states, both for static-structure comparisons (Table S2) and constant-pH simulations (Fig. 2B), we found the negatively charged (deprotonated) form of loop-F residue E160 was relatively preferred in the open versus closed states. At pH 6.5, the closed state is associated with a predominantly neutral (protonation fraction 0.55) form of E160, which formed transient hydrogen bonds with atoms including the backbone nitrogen of I20 on the same face (Fig. 3A-B). Conversely, the open state was associated with a negatively charged form of E160 (protonation fraction 0.1) (Fig. 3B). In this state, loop F rearranges to orient E160 away from I20 and the intrasubunit pocket; instead, E160 faced the relatively solvated interface between the extracellular/transmembrane domains and principal/complementary subunits, consistent with its solution-like p*K*_a_. Under alkaline conditions corresponding to maximal channel activity (pH 10), E160 further interacted with sodium ions accumulating in an electronegative cluster around the deprotonated side chains of E28 and E159 (Fig. 3A-B). These sodium ions frequently translocated in and out of the channel vestibule, indicating a lateral conduction pathway to and from the pore (Fig. S3). Thus the deprotonated form of E160 favors interactions in the open state, including an attraction for permeant ions, and could contribute to channel activity under alkaline conditions.

**Figure 3:**
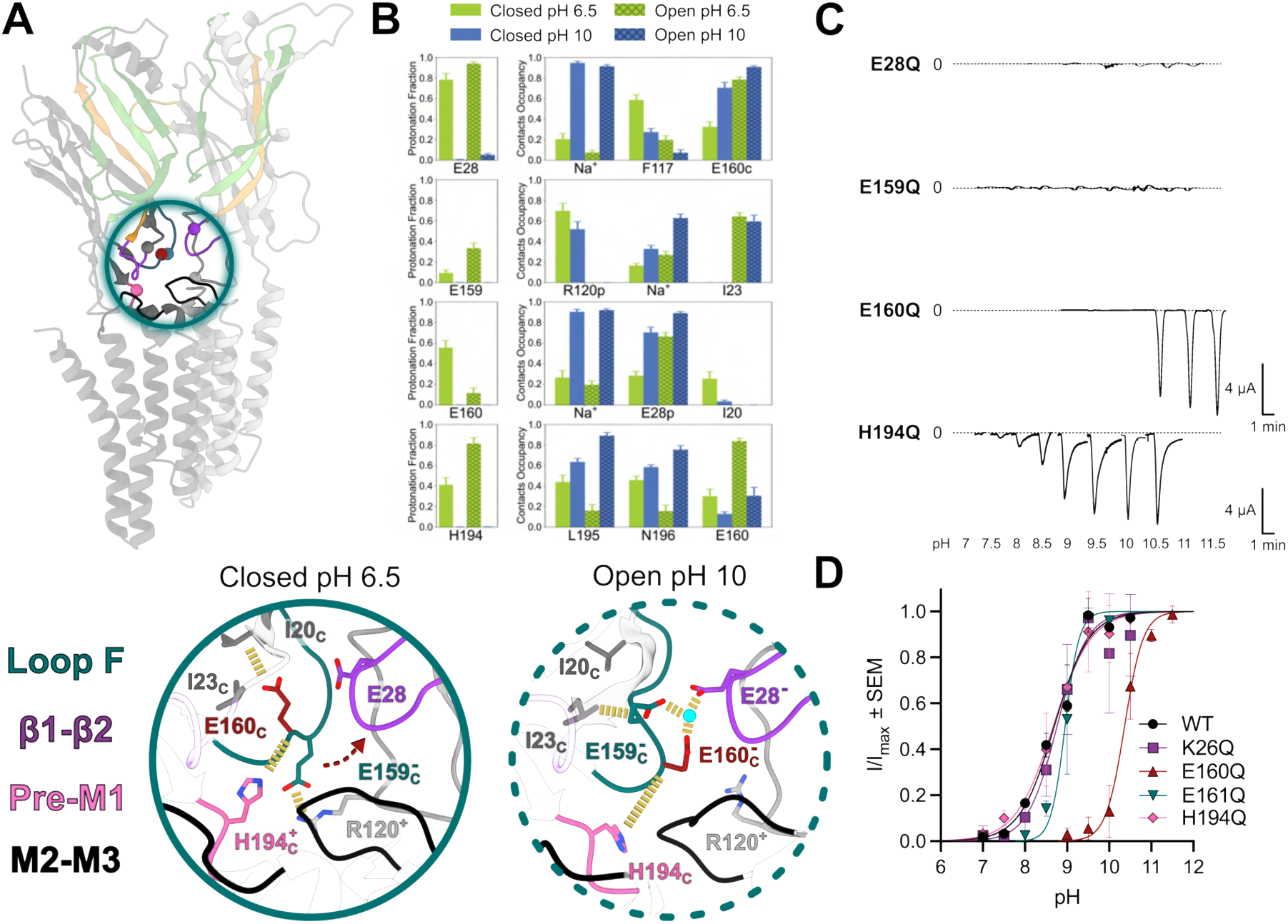
Deprotonation at E160 promotes activation and Na^+^ accumulation. **(A, top)** Two adjacent subunits are shown (PDB ID 6EX9 [16]), highlighting the location of titratable residues and their contact partners (spheres) at the ECD-TMD interface. **(A, bottom)** Schematics of the circled region in panel A highlighting differential interactions in the closed state at pH 6.5 (solid) and the open state at pH 10 (dotted) (PDB ID 9EWL [16]). Key side chains, including E160 (red), are represented as sticks and labeled. Ribbons and side chains associated with the ω1-ω2 loop, ω8-ω9 loop (loop F), pre-M1 region, and M2-M3 loop are colored purple, teal, pink, and black, respectively. Yellow dashes indicate state-specific polar contacts. **(B)** Mean protonation fraction and hydrogen-bond occupancies of contact partners for E28, E159, E160, and H194.**(C)** Representative oocyte electrophysiology traces for E28Q, E159Q, E160Q, and H194Q. **(D)** pH-response curves for WT, K26Q, E160Q, E161Q, and H194Q (n = 5). Normalized currents (I/I*_max_*) *±* SEM were fitted by nonlinear regression.

In order to test this hypothesis, Glu at position 160 was mutated to Gln in a sTeLIC construct appropriate for expression in *Xenopus* oocytes [15]. We predicted that substituting an amide for the carboxylate moiety would effectively lock the side chain in its neutral form, without substantially altering geometry or other physicochemical properties. If deprotonation of E160 is indeed a key contributor to alkaline activation, keeping this side chain neutral should favor the closed versus open state, and shift the activation curve to higher pH values. As previously reported by our group and others [16], wild-type sTeLIC in oocytes conducted 50 % maximal currents at pH 8.7 ± 0.1. The E160Q variant was substantially less pH sensitive, achieving 50% maximal activation only at pH 10.4 ± 0.1 (Fig. 3C-D and Table S1). Thus, electrophysiology recordings were consistent with deprotonation of E160 playing a key role in channel activation.

State dependence was also apparent, in both static-structure comparisons (Table S2) and constant-pH simulations (Fig. 2B), at K26 under alkaline conditions (pH 10). This basic residue is located at the same domain interface as E160, but on the ω1–ω2 loop of the principal subunit (Fig. S4). In the closed state, positively charged (protonated) K26 formed a persistent intersubunit salt bridge with E161, adjacent to E160 on complementary loop F (Fig. S4, S5A). Opening rearrangements in this region oriented K26 and E161 away from each other, instead contacting the M2–M3 loop of the principal-subunit TMD (residues S245 and R249 respectively). However, the open state was associated with only a modest loss of protonation at K26 (fraction 0.95 to 0.8), and E161 was primarily deprotonated throughout simulations in all conditions (Fig. S5A). Therefore, we predicted that substituting Gln at K26 or E161 would have limited effects on pH activation. Indeed, electrophysiology recordings showed pH sensitivities not significantly different from wild-type in both K26Q and E161Q variants (Fig. 3C-D, Fig. S6, and Table S1), consistent with these residues being relatively unimportant for sTeLIC gating.

At other residues in this region, evidence for state dependence was more complex. A His on the pre-M1 loop proximal to loop F (H194) was predicted by static-structure analysis to prefer double protonation, making it positively charged in the closed state at pH 6.5 (Table S2). However, local dynamics in our simulations inverted this preference, with greater double protonation in the open state (Fig. 3B). This simulated propensity contrasts with the anticipated behavior of residues critical to alkaline gating, as increasing pH would decrease protonation and favor a closed-rather than open-state environment for this residue. Indeed, our electrophysiology recordings indicated no significant effect of an H194Q substitution on pH sensitivity (Fig. 3C-D); its state-dependent shifts in protonation may reflect structural consequences of conformational change driven by e.g. E160, rather than a key role in activation.

Acidic residues proximal to E160 also exhibited complex behavior. Similar to H194, a Glu on the ω2 strand (E28) was predicted to prefer protonation in the closed state based on static-structure analysis (Table S2), but was protonated more extensively in the open state based on our constant-pH simulations (Fig. 3B). Specifically, E28 was already largely neutral (protonated) in the closed state, forming frequent intrasubunit hydrogen bonds with the backbone of F117; its modest increase in protonation in the open state appeared to accompany the increased proximity of the electronegative side chain of E160 (Fig. 3A-B). Similarly, a Glu immediately prior to E160 in sequence (E159) preferred protonation in the open state (Fig. 3B). The closed state was associated with an intersubunit salt bridge with R120 that could keep the side chain of E159 negative (deprotonated); in the open state, E159 formed largely intrasubunit polar interactions with the backbone of I23, creating less of a charge bias. Like H194, these patterns suggested that E28 and E159 do not critically promote sTeLIC gating, but are rather influenced by changes in the local environment during activation. On the other hand, both E28 and E159 contribute to the apparent accumulation site for sodium ions at the subunit interface, at least in the open state at pH 10, where both were fully deprotonated (Fig. 3B). Interestingly, multiple attempts to express E28Q and E159Q variants in oocytes yielded no consistent alkaline-gated currents (Fig. 3C), supporting a role for these residues in permeation or biogenesis independent of pH gating.

### Deprotonation of E106 in the Vestibule is Linked to ECD Contraction

Similar to E160, our simulations indicated a state-dependent shift in protonation at ω6 residue E106, for which the negatively charged (deprotonated) form is relatively preferred in the open versus closed states (Fig. 4A-B). Interestingly, static-structure analysis only predicted a 5 % shift in protonation at this position (Table S2), while our simulations captured a shift of 22% (Fig. 4B). E160 lies in a pocket surrounded by the ω4–ω5 strands (Λ-loop) within each subunit, facing the ECD vestibule. Channel activation has been shown to involve an overall contraction of the ECD, largely mediated by rigid-body motions of individual subunit ECDs towards their neighbors and the vestibule, increasing intersubunit contacts between neighboring Λ-loops. This transition has been implicated in positive allosteric modulation by psychostimulants in both sTeLIC and mammalian homologs, including direct drug interactions with ω6-Y104 [16] (Fig. S7). Interestingly, E106 formed consistent polar contacts with ω6-Y104 under all our simulated conditions, suggesting it can also contribute to relevant interaction networks (Fig. 4B, and Fig. S5B).

**Figure 4:**
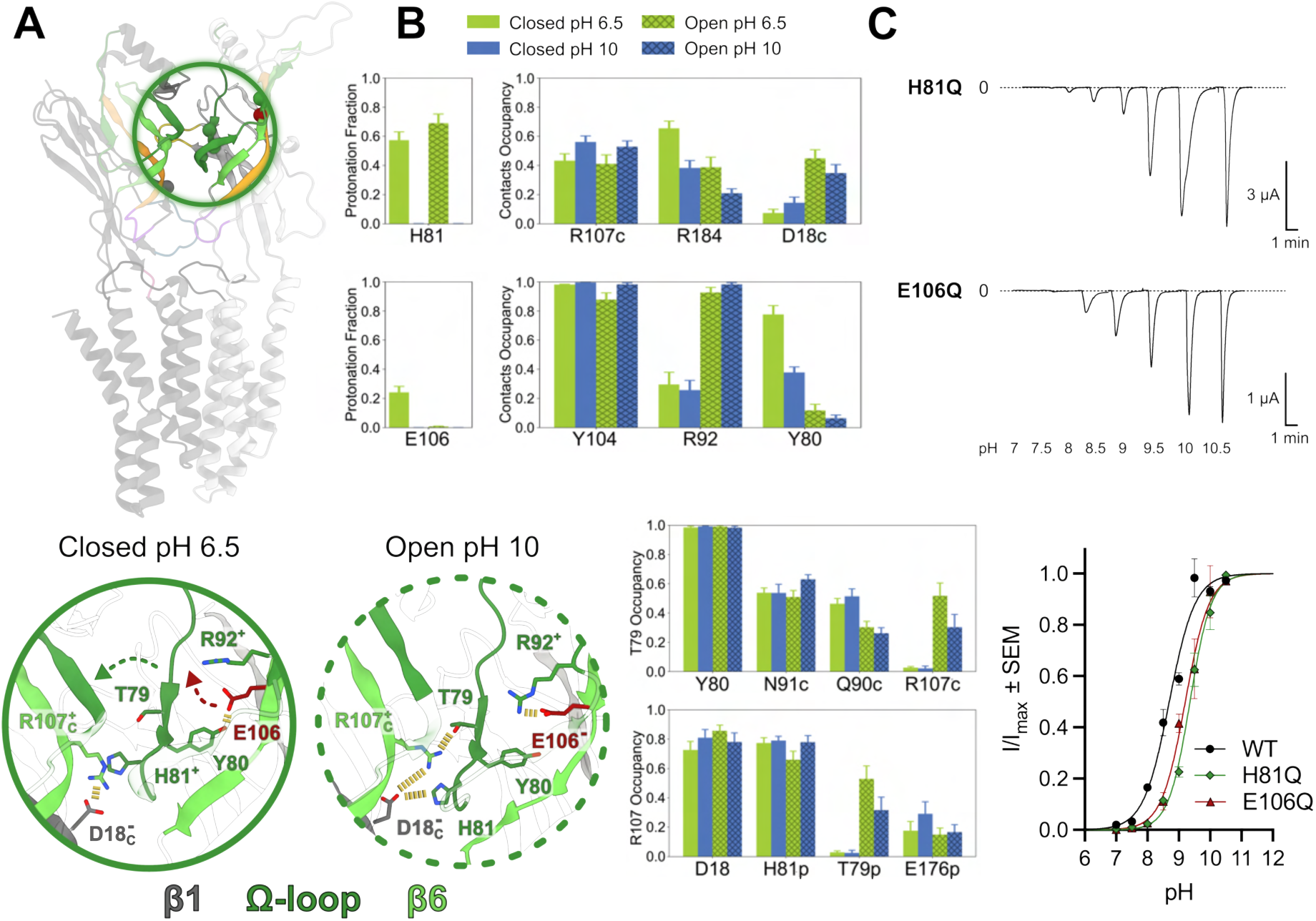
Deprotonation of E106 in the vestibule promotes open-state ECD contraction. **(A, top)** Two adjacent subunits are shown (PDB ID 6EX9 [16]), highlighting the location of titratable residues and their contact partners (spheres) within the vestibule.**(A, bottom)** Schematics of the circled region in panel A highlighting differential interactions in the closed state at pH 6.5 (solid) and the open state at pH 10 (dotted) (PDB ID 9EWL [16]). Key side chains, including E106 (red), are represented as sticks and labeled. Ribbons and side chains associated with the ω1, ω4-ω5 (!-loop), and ω6 strands are colored olive, dark, and light green, respectively. Yellow dashes indicate state-specific polar contacts. **(B)** Hydrogen-bond occupancies of the most frequent contact partners for ω4-H81, ω6-E106, ω4-T79, and ω6-R107. Columns represent simulations of the closed (solid) and open (crossed) states at pH 6.5 (lime) and pH 10 (blue), with error bars representing standard error of the mean. For ease of reference, mean protonation fractions are also reproduced from Fig. 2B for H81 and E106. **(C, top)** Representative oocyte electrophysiology traces for H81Q, and E106Q. **(C, bottom)** pH-response curves for WT, H81Q, and E106Q (n = 5). Normalized currents (I/I*_max_*) *±* SEM were fitted by nonlinear regression.

In the closed state, E106 was partially neutral (protonation fraction 0.25), and largely participated in intrasubunit polar interactions with ω4-Y80 in addition to ω6-Y104 (Fig. 4A-B, Fig. S5B). In the open state, E106 fully deprotonated, and shifted its interaction to the protonated ω5-R92 (Fig. 4A-B). Released from ω6-E106, ω4 interactions shifted toward the complementary subunit, as observed via increased intersubunit contacts of ω4-T79 and ω4-H81 (on either side of Y80) with residues including ω6-R107 and ω2-D18, respectively. Thus, our simulations provide dynamic details of vestibular rearrangements known to influence channel opening, linking deprotonation of ω6-E106 to the reorientation of multiple residues on ω4 towards the neighboring subunit.

Given the apparent contributions of both the ω4 and ω6 strands to pH-sensitive gating, we tested Gln mutations at H81 and E106 in oocytes. Substituting Gln at E106 neutralizes the side chain without substantially altering its geometry; we predicted this would favor closed-state interactions with ω4-Y80 over open-state interactions with ω5-R92, and shift the activation curve to higher pH values. Although H81 was not predicted as a major pH sensor in our simulations, substituting Gln at this position would reduce the side-chain volume and rigidity, as well as prevent its double protonation; we predicted these changes could limit open-state intersubunit contacts with D18, and again right-shift the pH activation curve. Indeed, activation curves for both the H81Q and E106Q variants were right-shifted by 0.5 pH units (Fig. 4C, and Table S1). Thus, simulations and recordings supported a role for the extracellular vestibule in alkaline pH gating, albeit with lesser influence than the domain-interface region surrounding E160.

Channel opening also increased intersubunit interactions further along the ω4 strand, for instance enabling W75 to contact the complementary ω3-T58 (Fig. S7A-B). W75 has been previously characterized as a key determinant of channel activation, flipping from intra- to intersubunit orientations upon opening [16]. Indeed, our simulations showed W75 predominantly faced the vestibular pocket within a single subunit in the closed state, but sampled an alternative rotamer toward the intersubunit interface in the open state (Fig. S7A, and C). Moreover, time-dependent plots of W75 showed reversible flipping between intra- and intersubunit orientations: the intersubunit rotamer was sampled only transiently in the closed state, but for prolonged periods in the open state (Fig. S8). Interestingly, the intersubunit rotamer never appeared in all five subunits simultaneously, even in open-state simulations. We previously found that psychostimulant binding in all five subunits forces W75 out of the vestibular pocket and into the intersubunit orientation [16]. Thus, our simulations are consistent with a vestibular mechanism of promoting activation, primed for further enhancement by ligand binding.

### Orthosteric Site Plays a Minimal Role in pH Gating

Centered at the ECD subunit, the so-called orthosteric site plays an essential role in eukaryotic pLGIC agonist binding [6]. However, we observed little rearrangement in this region between closed and open sTeLIC structures. In our simulations, channel opening relieves a modest neutralization of D39 (protonation fraction 0.21 to 0.04) (Fig. 2B). Similarly, static-structure calculations (though not our simulations) predicted a decrease in double protonation at H174 in the open state (Table S2). However, interaction patterns of these residues were not noticeably state-dependent (Fig. 5A-B). Other titratable residues in this region showed negligible (R37, H56, E176) or inverse (K179) pH sensitivity, and little evidence of state-dependent rearrangements (Figs. 5A-B, and Fig. S5C). To confirm the relative unimportance of this region in pH gating, we substituted Gln at D39, H174, E176, and K179; indeed, none of these variants exhibited a substantial shift in pH sensitivity (Fig. 5C). Thus, the domain-interface and extracellular-vestibule regions appear to contribute more than the orthosteric site to pH gating in sTeLIC.

**Figure 5:**
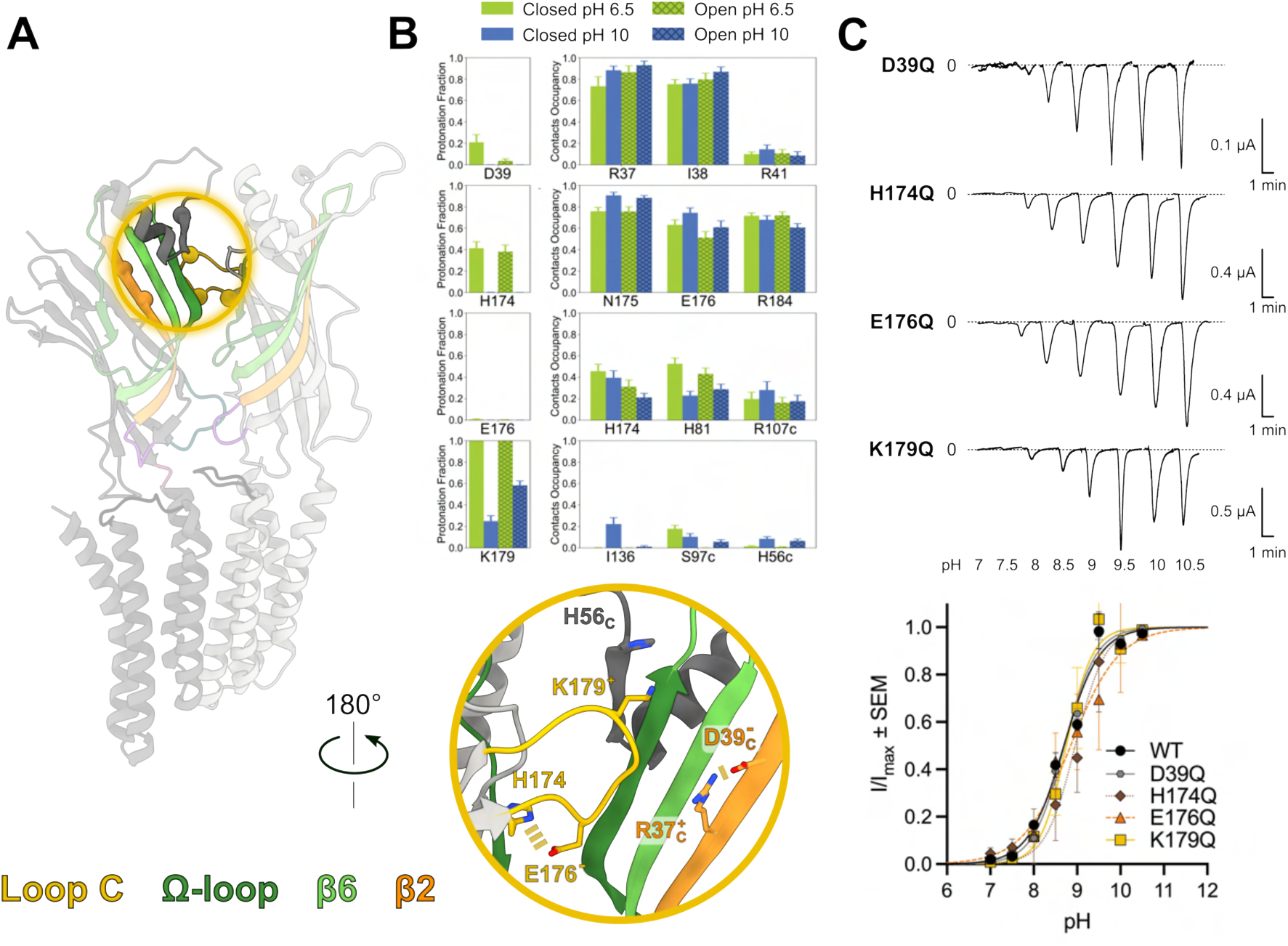
Orthosteric site plays a minimal role in pH gating. **(A, top)** Two adjacent subunits are shown (PDB ID 6EX9 [16]), highlighting the location of titratable residues and their contact partners (spheres) around the orthosteric site. **(A, bottom)** Schematic of the circled region in panel A providing a snapshot of consistent contact patterns in the closed and the open states. Potentially titratable side chains are represented as sticks and labeled. Ribbons and side chains associated with ω2, ω5 (!-loop), ω6, and the ω9-ω10 loop (loop C) are colored orange, dark green, light green, and yellow, respectively. **(B)** Mean protonation fraction and hydrogen-bond occupancies of contact partners for D39, H174, E176, and K179. Columns represent simulations of the closed (solid) and open (crossed) states at both pH 6.5 (lime) and pH 10 (blue), with error bars representing standard error of the mean. **(C, top)** Representative oocyte electrophysiology traces for D39, H174, E176, and K179. **(C, bottom)** pH-response curves for WT, D39, H174, E176, and K179 (n = 5). Normalized currents (I/I*_max_*) *±* SEM were fitted by nonlinear regression.

## Discussion

In this work, constant-pH MD simulations and electrophysiology recordings allowed us to discover titratable residues critical to pH gating in the model pLGIC sTeLIC, and to propose their mechanistic effects. In a plausible mechanistic model, deprotonation of acidic residues in both the domain interface (E160) and extracellular vestibule (E106) enables charge interactions associated with the open state, promoting side-chain remodeling particularly in the loop F and the Λ-loop (Fig. 6A). The domain-interface region also plays a role in permeating ions, while the extracellular vestibule is additionally involved in drug potentiation; thus, pH sensing appears to be distributed among mechanistically important elements of the receptor. Whereas some critical factors (e.g. E160) were anticipated by static-structure calculations, others (e.g. E106) were evident only in our simulations, demonstrating the importance of dynamics as well as functional validation in capturing relevant interactions. This work elaborates on previous constant-pH MD studies, for example of acid-sensing residues distributed across the ECD of GLIC (Fig. 6B) [22], with the additional consideration of residues titratable at alkaline pH. Despite our expectation that alkaline gating would be mediated by relatively basic amino acids, the most critical pH sensors we identified were acidic Glu residues, and it will be interesting to observe whether pH sensitivity in other ion channels follows similar patterns.

**Figure 6:**
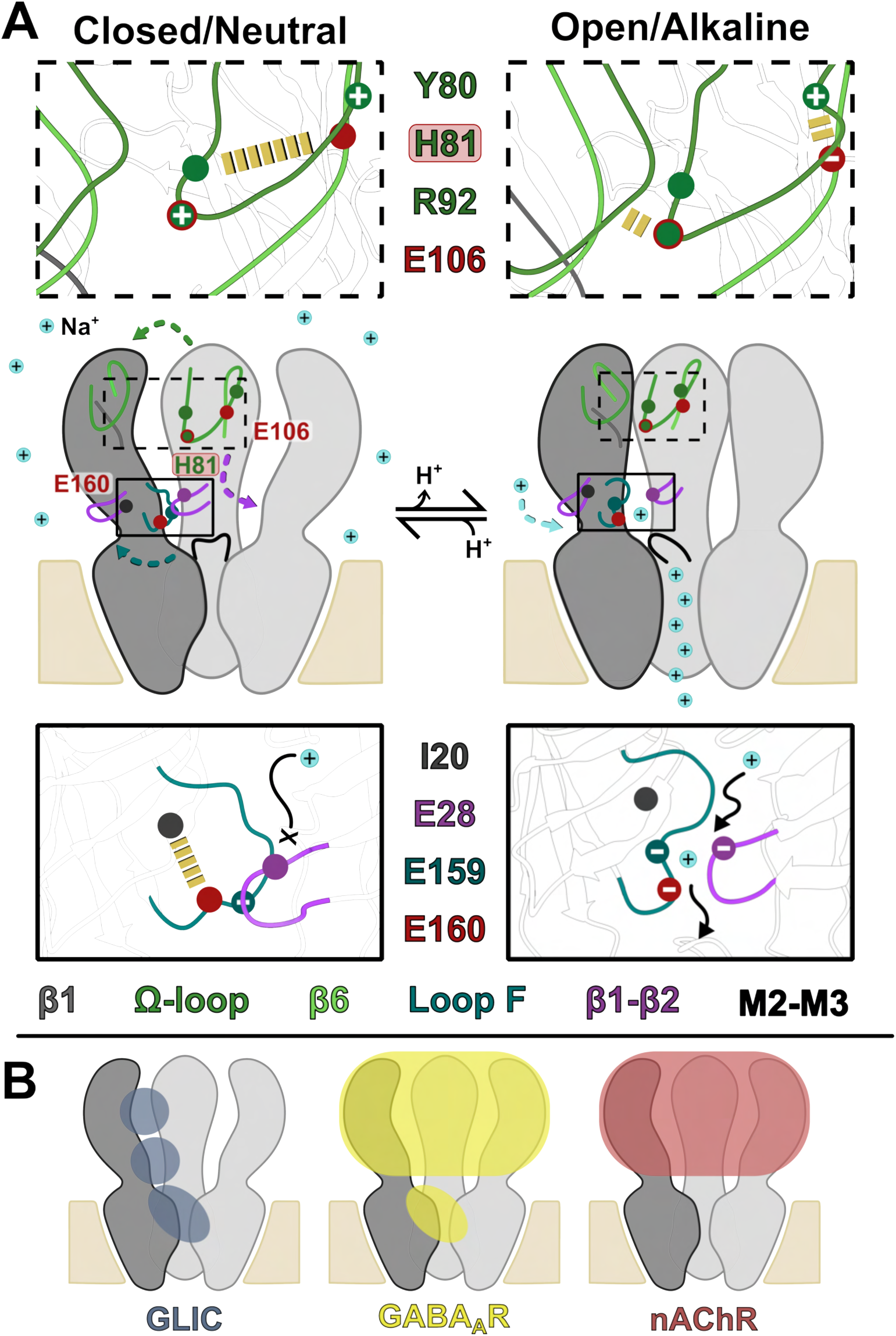
Proposed mechanism and generalizability of sTeLIC pH gating. **(A)** Schematic of sTeLIC with three adjacent subunits viewed from the membrane plane (side). In the proposed mechanism, the closed state at neutral pH (left) contains predominantly polar intrasubunit contacts of E106 and E160 (red) with their respective interaction partners (yellow dashes), both in the extracellular vestibule (upper inset) and domain interface (lower inset). Alkaline pH (right) promotes deprotonation of these Glu residues, weakening their closed-state contacts in favor of charge interactions. In the vestibule, this rearrangement facilitates intersubunit contacts and overall contraction between ECD subunits. At the domain interface, it facilitates dilation of the upper pore and permeation of positive ions (cyan spheres). In addition to E106 and E160, neighboring residues in both regions (colored spheres) may contribute directly to pH sensing and/or indirectly to channel activation. **(B)** Regions previously implicated in pH sensing in GLIC (blue), ϑ-aminobutyric acid-type A receptors (GABA_A_Rs, yellow), and nicotinic acetylcholine receptors (nAChRs, red).

In both our constant-pH and electrophysiology experiments, the principal mediator of sTeLIC pH gating appears to be E160 in loop F. Peripherally associated with the orthosteric site, loop F has long been implicated in pLGIC gating, but its mechanistic role remains unclear [25]. E160 exhibits dramatic deprotonation in the open versus closed state at equivalent pH, while protonation at adjacent titratable residues is relatively insensitive or inversely sensitive to activation. Deprotonation of E160 appears to weaken its intrasubunit hydrogen bond with I20 in favor of solvation at the domain/subunit interface in an acidic cluster with E28 and E159; the associated rearrangement of loop F is associated with outward dilation of the M2-M3 loops and, more distally, of the TMD pore. Interestingly, residues corresponding to E28 and E160 in GLIC (E35 and T158) have been implicated in that channel’s acidic gating mechanism, forming an intersubunit contact specifically in the open state [26]. Thus, the location and interactions involved in pH sensing may be at least partly conserved across evolution, even when the direction of pH sensitivity is reversed.

In its open-state orientation, E160 further contributes with E28 and E159 to Na^+^ ion interactions. Even though our simulations were conducted without a driving force, cation binding in this region is likely a key factor in their accumulation and permeation in and out of the channel. Access of Na^+^ ions via the domain/subunit interface is consistent with previous crystallographic studies of sTeLIC, which demonstrated Cs^+^ binding proximal to the E28-E159-E160 cluster, and predicted a lateral pathway for ions to and from the pore [15]. Aside from its evident role in ion permeation in a number of pLGICs, this interface has also been shown to bind Ca^2+^ in ELIC [27], DeCLIC [14], and nicotinic acetylcholine receptors (nAChRs) [28], as well as Zn^2+^ in type-A ϑ-aminobutyric acid receptors (GABA_A_Rs) [29]. Although the structural basis for sTeLIC inhibition by Ca^2+^ was beyond the scope of this work, it is plausible that the region around loop F constitutes a convergent site for both ion permeation and ion block.

The extracellular vestibule also influences sTeLIC pH gating, via direct and/or indirect effects involving H81 and E106. We recently showed the vestibular site within each sTeLIC subunit binds psychostimulants, particularly via residues W75 and Y104, thereby facilitating channel activation [16]. In that work, the mechanism for sTeLIC activation in the absence of drug remained ambiguous, since no open experimental structure has been captured so far without evident ligand density in the vestibular pocket. Here, we find that deprotonation of E106 (proximal to Y104 in the drug pocket) weakens its intrasubunit hydrogen bond with ω4-Y80 in favor of a charge interaction with ω5-R92, on the opposite side of the Λ-loop. Adjacent to Y80, ω4-H81 is accordingly able to form more frequent contacts with the neighboring subunit, consistent with open-state contraction of the ECD. Mutations at these positions produced shifted pH sensitivity, albeit more modestly than at E160, indicating they make a secondary contribution to the alkaline gating mechanism. Other ω4 residues including W75 also increase intersubunit contacts, consistent with their involvement in channel activation, and are poised for further enhancement upon drug binding. Thus, our simulations offer insights into the role of the extracellular vestibule in both pH gating and drug potentiation.

Despite its well-documented role in eukaryotic pLGIC gating, the orthosteric site did not substantially influence alkaline gating in sTeLIC. Titratable residues in this region exhibited little rearrangement in their interaction patterns between closed and open simulations, and failed to substantially alter pH sensitivity when mutated. This observation echoes the related channel GLIC, where loop C has been shown to play little role in acidic-pH [30]. Compared to ligand gating, pH gating appears to arise by differential, more distributed evolutionary pressures involving protonation changes elsewhere in the ECD. Indeed, pH sensitivity has been reported parallel to ligand gating in eukaryotic pLGICs gated by GABA [8–10, 31], glycine [12], and acetylcholine [11], involving multiple potential sites in the ECD and upper TMD (Fig. 6B); our present work offers a template for more detailed characterization of the structural basis for such effects.

## Methods

### Simulation Protocols

#### System Setup

Closed (PDB ID 9EX6) and open (PDB ID 9EWL) structures [16] of sTeLIC were used as starting models for constant-pH MD simulations. Each structure was placed in a 13.0 nm *→* 13.0 nm *→* 15.6 nm box and embedded in a bilayer formed by 1-palmitoyl-2-oleoylphosphatidylcholine (POPC) lipids (408 for the closed state, 417 for the open state) using the CHARMM-GUI membrane builder [32]. The CHARMM36-mar2019-cphmd force field [33] and the CHARMM TIP3P water model were applied during topology generation. The term *cphmd* indicates that modifications specific to constant-pH MD have been incorporated into the bonded parameters for the titratable amino acids Asp, Glu, His, and Lys. Further information regarding these modifications can be found in [34]. The simulation input files were established through phbuilder [35]. Initially, all Asp, Glu, His, Arg, Tyr, and Lys residues were made titratable. The constant-pH MD parameters for Asp, Glu, Lys, and His were acquired from [34], while those for Arg were obtained from [35]. For Tyr, the partial charges on the titratable atoms in the deprotonated state were calculated using the CHARMM-GUI Ligand Modeler [36], and the resulting charges are provided in Table S3. The parameterization of the correction potential was conducted in accordance with the methodology outlined in [35], while the reference p*K*_a_ was obtained from [21]. The finalized correction potential is included in the associated Zenodo deposition. Upon the selection of the titratable residues, NaCl was added to ensure a net-neutral environment at *t* = 0 and to set an ion concentration of 150 mM. Furthermore, 300 buffer particles were added to address charge fluctuations and maintain a net-neutral system for *t >* 0. For further details regarding the buffer particles, see [18, 37]. phbuilder was employed to create all constant-pH MD-specific GROMACS input parameters.

#### Constant-pH Molecular Dynamics Simulations

All constant-pH MD simulations were conducted using the GROMACS constant-pH MD beta version. This version is based on the 2021 release branch and is modified to include the routines required for performing the *ω*-dynamics calculations. The source code branch can be accessed at www.gitlab.com/gromacs-constantph until it is fully integrated into the main distribution.

Energy minimization was carried out using the steepest-descent algorithm. Initial relaxation was performed in the NVT ensemble for 10 ps, using a time step of 0.2 fs and the v-rescale thermostat [38], which operated with a coupling time of 0.5 ps at a temperature of 300 K. Lengths of bonds involving hydrogens were constrained through the P-LINCS algorithm [39], while electrostatic interactions were calculated using the PME method [40]. Following this, further relaxation and production runs were executed in the NPT ensemble with a time step of 2 fs, maintaining a pressure of 1 bar via the c-rescale barostat [41], which had a coupling time of 5 ps. After 1 ns of relaxation, the *ω*-coordinates were released while the heavy atoms were kept restrained. An additional relaxation phase of 30 ns followed, during which the restraints on the heavy atoms were gradually released.

To assess whether the default bias potential barriers of 7.5 kJ/mol adequately restricted the sampling of unphysical states where *ω ↑*= 0, 1, a series of 20 ns test runs were conducted. Several affected residues were identified,and the inverse Boltzmann method was used to calculate the necessary increase in barrier height for these residues. In the closed state simulations at pH 6.5, the residues D39 (20 kJ/mol) and D134 (15 kJ/mol) were found to be affected. At pH 10, both closed and open state simulations revealed that residues K71 (10 kJ/mol), K179 (10 kJ/mol), and Y300 (20 kJ/mol) were impacted. In the open state simulations at pH 6.5, only the residue D134 (10 kJ/mol) was affected.

#### Analysis

The mean protonation fractions are derived by averaging the fifteen time-averaged values of protonation fractions from the five subunits across three replicates. The protonation fractions are defined as *N*_proto_*/*(*N*_proto_ + *N*_deproto_), where *N*_proto_ and *N*_deproto_ represent the number of simulation frames in which the residue is classified as protonated (*ω <* 0.2) or deprotonated (*ω >* 0.8), respectively. Contact analysis of the titratable residues is conducted between the carboxyl oxygens (Asp, Glu), the ring nitrogens (His), the amino nitrogens (Arg, Lys), the hydroxyl oxygen (Tyr), and the polar atoms (oxygen, nitrogen) of contacting residues. A contact cutoff of 0.4 nm is applied to all interactions. Residues undergoing state-dependent protonation changes are selected based on the mean protonation fractions *±* SEM and evaluated using unpaired Student’s *t* tests, with a significance threshold set at P *<* 0.05. Hydration and pore properties are analyzed in CHAP [19], with additional plots of the 9’ radius (defined as the shortest distance from the closest atom in L234 of a given subunit to the C*_ω_*-center of mass of all L234 residues) produced in MDAnalysis [42].

### Experimental Protocols

#### Mutagenesis Design and Generation

Mutations generated in sTeLIC were inserted into the pMT3 expression vector, using the GeneArt Site-Directed Mutagenesis System (Thermo Fisher Scientific) and commercial mutagenic primers (IDT). The PCR products were digested overnight with DpnI at 37°C and then transformed into XL1-Blue Supercompetent cells (Agilent). Verification of mutant cDNAs was achieved by Sanger sequencing (Eurofins Genomics), followed by large-scale purification with a HiSpeed Plasmid Midi Kit (Qiagen).

#### Oocyte Preparation

Oocytes from *Xenopus laevis* frogs (Ecocyte Bioscience) were injected via the animal pole with 6 ng/32.2 nl sTeLIC WT or mutant cDNA using a Nanoject II microinjector (Drummond Scientific). After injection, oocytes were kept in modified Barth’s solution (88 mM NaCl, 1 mM KCl, 2.4 mM NaHCO_3_, 0.91 mM CaCl_2_, 0.82 mM MgSO_4_, 0.33 mM Ca(NO_3_)_2_, 10 mM HEPES, 0.5 mM theophylline, 0.1 mM G418, 17 mM streptomycin, 10,000 U/l penicillin and 2 mM sodium pyruvate, adjusted to pH 7.5) at 13°C.

#### Oocyte Electrophysiology

pH-evoked currents were recorded 4–7 days after cDNA injection using a two-electrode voltage-clamp (TEVC). The oocyte membrane was maintained at constant voltage by using two sharp recording needles (5 - 50 MΛ) filled with 3 M KCl, along with two bath electrodes that connected the ground stages and the bath chamber through 3 M KCl-agar bridges. The oocytes were continuously perfused with a running buffer (123 mM NaCl, 2 mM KCl and 2 mM MgSO_4_, and adjusted to pH 6.5) at a flow rate of approximately 1 mL/min. The oocytes were maintained at a holding potential of -40 mV using an OC-725C voltage clamp (Warner Instruments). Digidata 1440A (Molecular Devices) was used to sample and digitize data using Clampex (Axon Instruments). Currents were filtered at a frequency of 1 kHz and analyzed using Clampfit (Axon Instruments). After establishing a stable baseline, pH buffers were applied in increments of 0.5 (pH 7 - pH 11.5) for a duration of 30 s, followed by 8-min washout periods.

#### Statistics

Statistical analyzes of electrophysiology data were performed using Prism 10.0.3 (GraphPad Software). The data analyzed were derived from groups consisting of a minimum of five independent experiments. The results are presented as mean *±* SEM and were evaluated using unpaired Student’s *t* tests, with a significance threshold set at P *<* 0.05. The normalized proton activation curves were calculated using the equation; *Y* = (*R_basal_*+((*R_max_ ↓ R_basal_*)*/*(1 + 10^[^*^logEC^*^50^*^→logX^*^]^*^↑nH^* ))), where Y is the activation response, R*_max_* is the maximal response, R*_basal_* is the baseline, X is the proton concentration, EC_50_ is the pH level that provokes a half-maximum response, and n is the Hill coefficient.

### Data and Software Availability

The GROMACS constant-pH beta is available at www.gitlab.com/gromacs-constantph.

## Abbreviations

CpHMD: Constant-pH molecular dynamics
EC_50_: Half-maximal effective concentration
ECD: Extracellular domain
pLGIC: Pentameric ligand-gated ion channel
RMSD: Root-mean-squared deviation
SEM: Standard error of mean
TEVC: Two-electrode voltage-clamp
TMD: Transmembrane domain
WT: Wild type

## Acknowledgments

We thank S. Eriksson Lidbrink for assistance with W75 figures and analysis. E.K. was supported by a PhD fellowship from the Sven och Lilly Lawskis fond. R.J.H., E.L. and B.H. were supported by grants from the Swedish Research Council (grant nos. 2019-02433, 2019-04477, and 2021-05806), the BioExcel Center of Excellence (grant nos. H2020-INFRAEDI-02-2018-823830 and EuroHPC 101093290), the Knut and Alice Wallenberg Foundation, and the Swedish e-Science Research Centre. Computational resources were provided by SNIC (grant nos. 2022/3-40 and 2022/21-16) and NAISS (grant no. 2023/3-27).

## Author Contributions

Conceptualization: E.K., R.J.H.; methodology: E.K., A.J., B.H.; software: A.J., B.H.; investigation - electrophysiology: E.K., J.P.; investigation - simulations: I.Y., A.J.; investigation - p*K*_a_ prediction: J.P.; analysis and visualization: E.K., I.Y.; writing - original draft: E.K.; writing - review and editing: E.K., R.J.H., E.L., B.H.; supervision: R.J.H.; project administration: R.J.H.; funding acquisition: E.L., B.H.

