## Supplementary Information for "Mechanism of Alkaline Gating in a Pentameric Ion Channel"

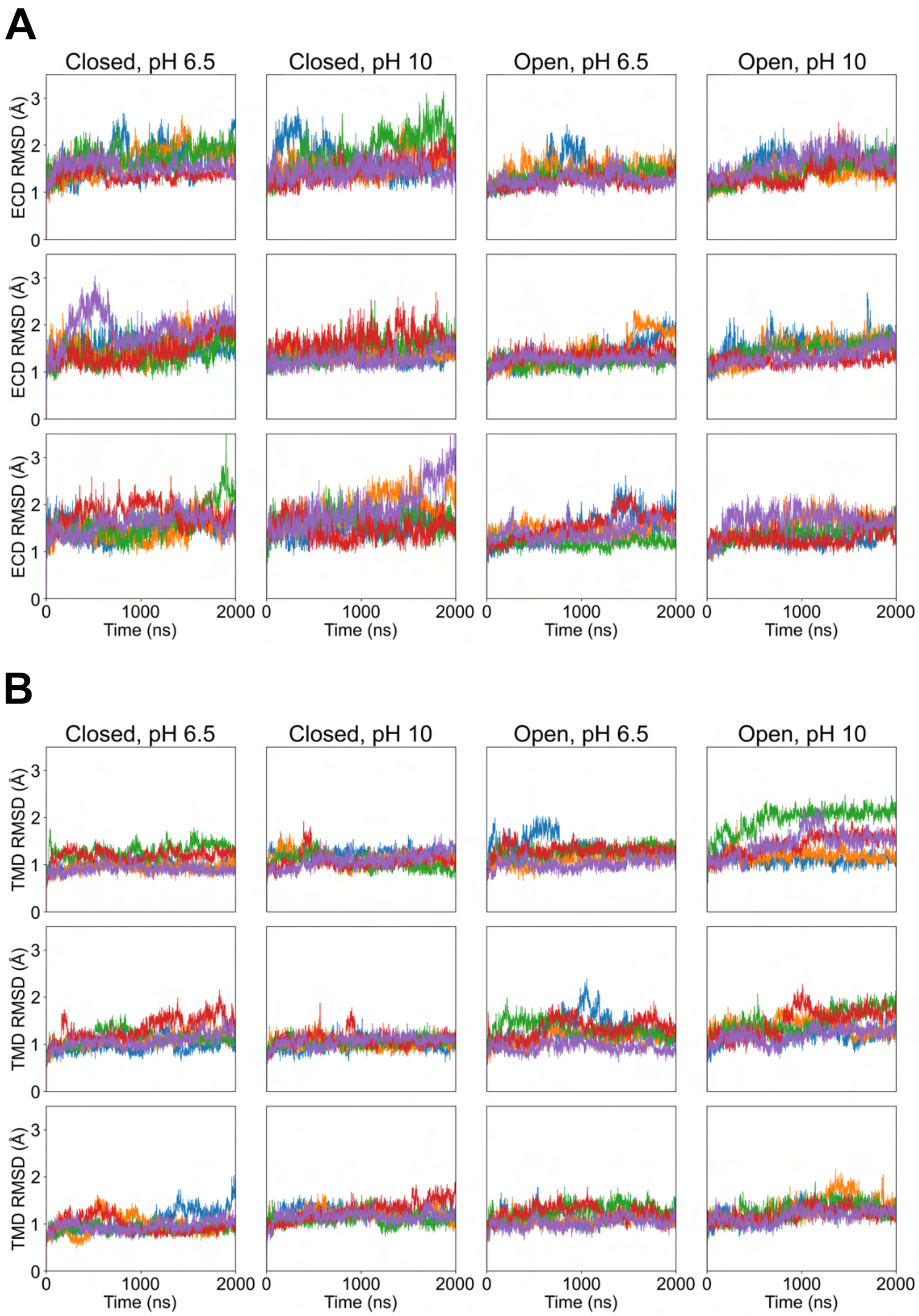


**Figure S1: Stability of sTeLIC domains in constant-pH MD simulations.** **(A)** $C_\alpha$ RMSD (Å) for residues 1–196 (ECD) in constant-pH simulations of the (left to right) closed structure at pH 6.5, closed structure at pH 10, open structure at pH 6.5, and open structure at pH 10. In each condition, the five sTeLIC subunits are represented as five separately colored plots. Three replicate simulations of each system are shown as separate rows. **(B)** $C_\alpha$ RMSD (Å) for residues 187–320 (TMD) in constant-pH simulations as in panel A.

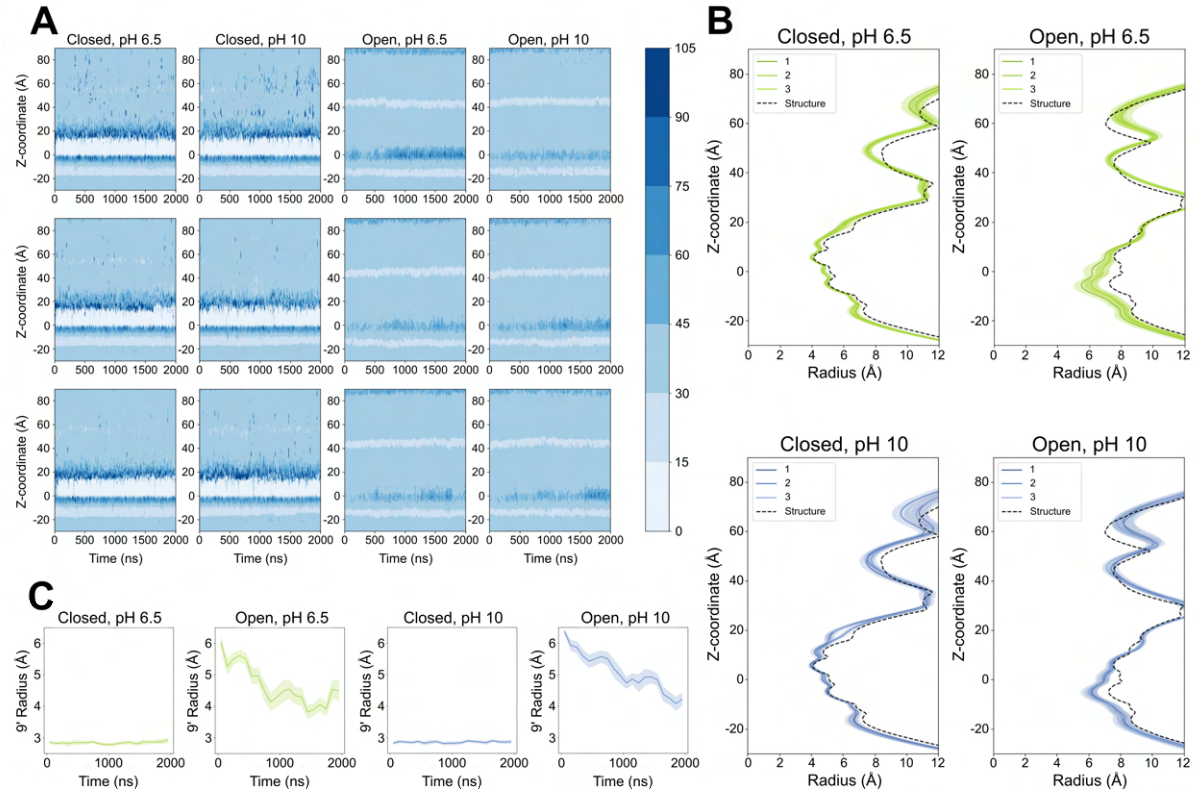


**Figure S2: Hydration and pore profiles.** **(A)** Water density in the pore in constant-pH simulations of the (left to right) closed structure at pH 6.5, closed structure at pH 10, open structure at pH 6.5, and open structure at pH 10, colored according to scalebar at right (molecules/nm$^3$), as calculated in CHAP [19]. Three replicate simulations of each system are shown as separate rows. **(B)** Backbone pore profiles, relative to the starting structure (dashed black line), for each replicate constant-pH simulation of the closed structure at pH 6.5 (top left, green), closed structure at pH 10 (lower left, blue), open structure at pH 6.5 (top right, green), and open structure at pH 10 (lower right, blue). Each profile is represented as mean (solid line) $\pm$ standard deviation (shaded). **(C)** Average radius at the 9' position across all replicate constant-pH simulations of the (left to right) closed structure at pH 6.5, open structure at pH 6.5, closed structure at pH 10, and open structure at pH 10. Each profile is represented as mean (solid line) $\pm$ standard deviation (shaded).

**Table S1: Simulated state dependence and electrophysiological pH sensitivity of mutants involving key sTeLIC residues.** P-value and significance correspond to the largest difference between mean protonation fractions for a given residue in closed- versus open-state simulations at the same pH, determined using unpaired Student's $t$ tests. Protonation fraction was considered significantly state dependent where $P \leq 0.05$ (*), $P \leq 0.01$ (**), $P \leq 0.001$ (***), or $P \leq 0.0001$ (****); $P > 0.05$ was presumed to indicate no significant difference (ns). $EC_{50}$ values are derived by nonlinear regression analysis of pH activation curves in at least 4 oocytes for variants with Gln substituted for the indicated residue. For residues labeled non-determined (ND), Gln-substituted variants did not produce measurable currents, even under the highest accessible pH conditions. *Shift* indicates the increase in $EC_{50}$ for a given variant with respect to wild-type sTeLIC ($EC_{50} = 8.7 \pm 0.1$, $n = 4$).

| **Residue** | **P-value** | **Significance** | **Variant $EC_{50}$** | **Shift** |
|---|---|---|---|---|
| K26 | < 0.0001 | **** | 8.7 ± 0.1 | 0.0 |
| E28 | 0.0291 | * | ND | ND |
| D39 | 0.0324 | * | 8.7 ± 0.1 | 0.0 |
| H81 | 0.1815 | ns | 9.3 ± 0.1 | 0.6 |
| E106 | < 0.0001 | **** | 9.2 ± 0.1 | 0.5 |
| E159 | 0.0003 | *** | ND | ND |
| E160 | < 0.0001 | **** | 10.4 ± 0.1 | 1.7 |
| E161 | 0.026 | ns | 9.0 ± 0.1 | 0.3 |
| H174 | 0.7089 | ns | 8.9 ± 0.1 | 0.2 |
| E176 | 0.3626 | ns | 8.8 ± 0.1 | 0.1 |
| K179 | < 0.0001 | **** | 8.7 ± 0.1 | 0.0 |
| H194 | 0.0001 | *** | 8.7 ± 0.1 | 0.0 |

**Table S2: Protonation fractions predicted from static structures.** Protonation fractions predicted for the closed (PDB ID 9EX6 [16]) or modulator-bound open structure (PDB ID 9EWL [16]) at pH 6.5 or 10, based on the p$K_a$s estimated in Prop$K_a$ shown for each structure at far right[23, 24]. Residues with a maximum difference in mean protonation fractions of more than 15 percentage points between the open and closed states at the same pH are highlighted in gray.

| | Protonation Fractions | | | | Absolute p$K_a$ | |
|---|---|---|---|---|---|---|
| Residue | Closed, pH 6.5 | Closed, pH 10 | Open, pH 6.5 | Open, pH 10 | Closed | Open |
| E7 | 0.01 | 0.00 | 0.01 | 0.00 | 4.50 | 4.54 |
| D10 | 0.00 | 0.00 | 0.00 | 0.00 | 3.47 | 3.90 |
| K16 | 1.00 | 0.78 | 1.00 | 0.81 | 10.84 | 11.38 |
| D18 | 0.00 | 0.00 | 0.00 | 0.00 | 1.26 | -0.70 |
| K26 | 1.00 | 0.88 | 1.00 | 0.56 | 10.92 | 9.93 |
| E27 | 0.02 | 0.00 | 0.01 | 0.00 | 4.84 | 4.43 |
| E28 | 0.45 | 0.00 | 0.24 | 0.15 | 6.41 | 6.05 |
| R37 | 1.00 | 1.00 | 1.00 | 0.99 | 13.63 | 12.31 |
| D39 | 0.00 | 0.00 | 0.00 | 0.00 | 2.30 | 2.70 |
| R41 | 1.00 | 0.99 | 1.00 | 1.00 | 11.94 | 12.33 |
| E48 | 0.01 | 0.00 | 0.01 | 0.00 | 4.50 | 4.50 |
| H49 | 0.50 | 0.00 | 0.75 | 0.00 | 6.50 | 6.97 |
| E53 | 0.00 | 0.00 | 0.01 | 0.00 | 3.89 | 4.41 |
| K55 | 1.00 | 0.76 | 1.00 | 0.72 | 10.50 | 10.42 |
| H56 | 0.49 | 0.00 | 0.37 | 0.00 | 6.49 | 5.99 |
| R57 | 1.00 | 0.99 | 1.00 | 0.99 | 11.87 | 11.98 |
| Y59 | 1.00 | 0.55 | 1.00 | 0.49 | 10.09 | 9.99 |
| K66 | 1.00 | 0.76 | 1.00 | 0.71 | 10.50 | 10.40 |
| E69 | 0.01 | 0.00 | 0.00 | 0.00 | 4.50 | 2.85 |
| E70 | 0.01 | 0.00 | 0.00 | 0.00 | 4.57 | 3.81 |
| K71 | 1.00 | 0.70 | 1.00 | 0.72 | 10.36 | 10.42 |
| R74 | 1.00 | 0.99 | 1.00 | 0.92 | 11.87 | 11.66 |
| Y80 | 1.00 | 1.00 | 1.00 | 1.00 | 16.25 | 14.72 |
| H81 | 0.08 | 0.00 | 0.93 | 0.20 | 5.35 | 8.32 |
| R86 | 1.00 | 0.99 | 1.00 | 0.99 | 12.29 | 11.98 |
| D88 | 0.00 | 0.00 | 0.01 | 0.00 | 4.01 | 4.06 |
| R92 | 1.00 | 1.00 | 1.00 | 1.00 | 13.35 | 13.93 |
| E98 | 0.01 | 0.00 | 0.00 | 0.00 | 4.64 | 4.17 |
| D99 | 0.00 | 0.00 | 0.00 | 0.00 | 1.38 | 3.18 |
| Y104 | 1.00 | 1.00 | 1.00 | 0.99 | 13.62 | 12.27 |
| E106 | 0.05 | 0.00 | 0.00 | 0.00 | 5.24 | 2.73 |
| R107 | 1.00 | 0.98 | 1.00 | 0.94 | 11.83 | 11.64 |
| D118 | 0.01 | 0.00 | 0.00 | 0.00 | 4.33 | 3.42 |
| R120 | 1.00 | 0.99 | 1.00 | 0.99 | 12.01 | 12.32 |
| D125 | 0.00 | 0.00 | 0.00 | 0.00 | 1.50 | 1.62 |
| H132 | 0.00 | 0.00 | 0.29 | 0.00 | 0.88 | 4.41 |
| D134 | 0.00 | 0.00 | 0.00 | 0.00 | 1.64 | 1.96 |

*Continued on next page*

| | | | | | | |
|---|---|---|---|---|---|---|
| H140 | 0.38 | 0.00 | 0.45 | 0.00 | 6.29 | 6.42 |
| R143 | 1.00 | 1.00 | 1.00 | 1.00 | 12.36 | 12.37 |
| E146 | 0.00 | 0.00 | 0.01 | 0.00 | 4.12 | 4.23 |
| D155 | 0.00 | 0.00 | 0.00 | 0.00 | 3.94 | 3.86 |
| E159 | 0.00 | 0.00 | 0.04 | 0.00 | 1.64 | 4.88 |
| E160 | 0.67 | 0.00 | 0.01 | 0.00 | 6.80 | 4.30 |
| E161 | 0.03 | 0.00 | 0.00 | 0.00 | 4.83 | 3.34 |
| E166 | 0.00 | 0.00 | 0.00 | 0.00 | 4.12 | 3.78 |
| H170 | 0.50 | 0.00 | 0.48 | 0.00 | 6.50 | 6.46 |
| H174 | 0.98 | 0.02 | 0.51 | 0.00 | 8.24 | 6.39 |
| E176 | 0.00 | 0.00 | 0.00 | 0.00 | -5.82 | 2.72 |
| K179 | 1.00 | 0.57 | 1.00 | 0.76 | 10.12 | 10.50 |
| D181 | 0.00 | 0.00 | 0.00 | 0.00 | 3.22 | 3.16 |
| R184 | 1.00 | 1.00 | 1.00 | 1.00 | 14.72 | 12.82 |
| E188 | 0.01 | 0.00 | 0.02 | 0.00 | 4.48 | 4.70 |
| H190 | 0.46 | 0.00 | 0.48 | 0.00 | 6.43 | 6.46 |
| E192 | 0.01 | 0.00 | 0.01 | 0.00 | 4.20 | 4.53 |
| R193 | 1.00 | 1.00 | 1.00 | 1.00 | 14.37 | 13.79 |
| H194 | 0.89 | 0.00 | 0.40 | 0.00 | 7.39 | 6.14 |
| Y197 | 1.00 | 1.00 | 1.00 | 0.79 | 14.99 | 10.97 |
| Y198 | 1.00 | 0.93 | 1.00 | 0.99 | 11.15 | 12.72 |
| R201 | 1.00 | 1.00 | 1.00 | 0.99 | 13.86 | 12.32 |
| D221 | 0.00 | 0.00 | 0.00 | 0.00 | 2.80 | 3.44 |
| Y222 | 1.00 | 0.89 | 1.00 | 1.00 | 10.93 | 13.17 |
| K224 | 1.00 | 0.54 | 1.00 | 0.72 | 10.35 | 10.50 |
| R225 | 1.00 | 1.00 | 1.00 | 0.96 | 13.41 | 12.27 |
| D227 | 0.00 | 0.00 | 0.00 | 0.00 | 3.96 | 2.76 |
| D246 | 0.00 | 0.00 | 0.00 | 0.00 | 3.13 | 2.12 |
| R249 | 1.00 | 0.98 | 1.00 | 1.00 | 11.67 | 14.97 |
| Y252 | 1.00 | 0.98 | 1.00 | 1.00 | 11.75 | 13.57 |
| D257 | 0.01 | 0.00 | 0.00 | 0.00 | 4.28 | 1.97 |
| R278 | 1.00 | 0.99 | 1.00 | 0.99 | 12.17 | 12.30 |
| R279 | 1.00 | 0.99 | 1.00 | 0.99 | 12.22 | 12.23 |
| E281 | 0.00 | 0.00 | 0.00 | 0.00 | 3.98 | 3.69 |
| H283 | 0.91 | 0.00 | 0.64 | 0.00 | 7.53 | 6.82 |
| K285 | 1.00 | 0.76 | 1.00 | 0.76 | 10.50 | 10.50 |
| R290 | 1.00 | 1.00 | 1.00 | 0.99 | 12.50 | 12.23 |
| K291 | 1.00 | 0.73 | 1.00 | 0.75 | 10.43 | 10.47 |
| D293 | 0.00 | 0.00 | 0.00 | 0.00 | 0.27 | 2.80 |
| Y295 | 1.00 | 0.54 | 1.00 | 0.50 | 10.07 | 10.00 |
| Y300 | 1.00 | 0.64 | 1.00 | 0.79 | 10.25 | 11.75 |
| Y304 | 1.00 | 1.00 | 1.00 | 0.31 | 14.55 | 9.47 |

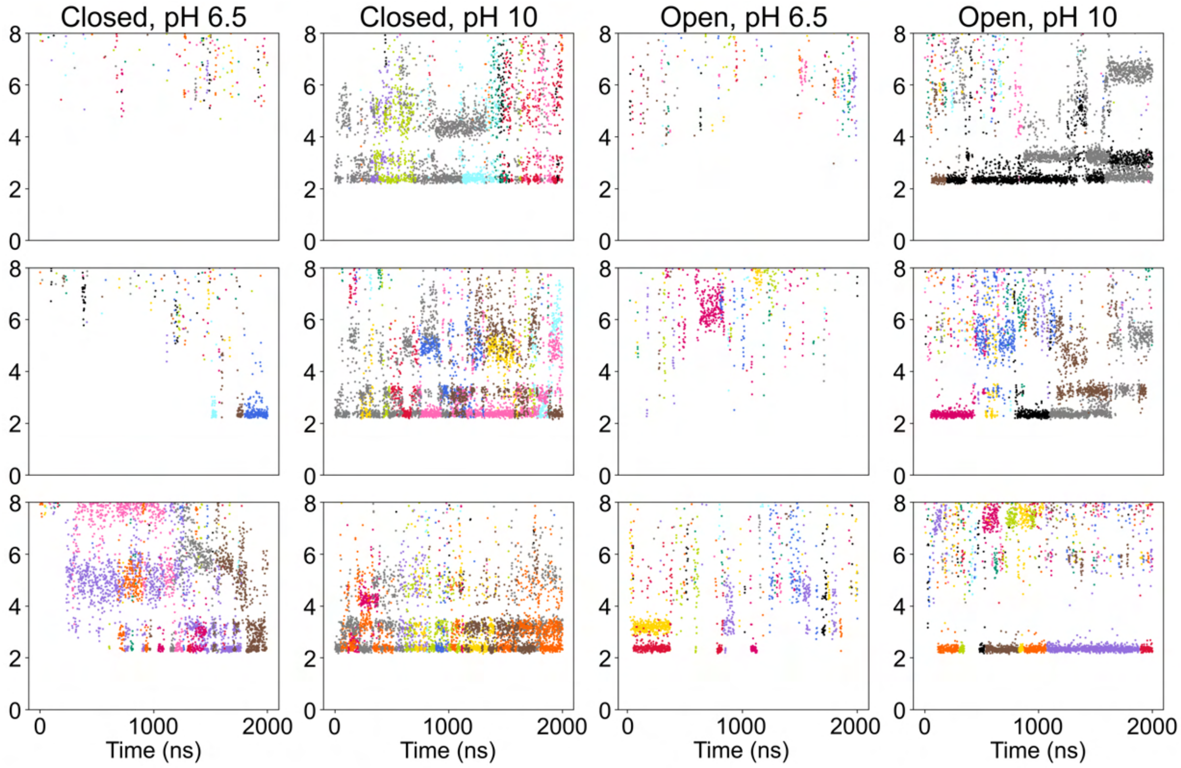


**Figure S3: Time-dependent $Na^+$ exchange in the interfacial pocket.** Each plot shows displacement of individual $Na^+$ ions, colored separately, from the terminal C atom of E28 residues at the five subunit/domain interfaces in a single constant-pH MD trajectory. Replicate simulations are represented in parallel rows. Under activating conditions (pH 10), ions accumulate in multiple positions $\geq 2$ Å from E28.

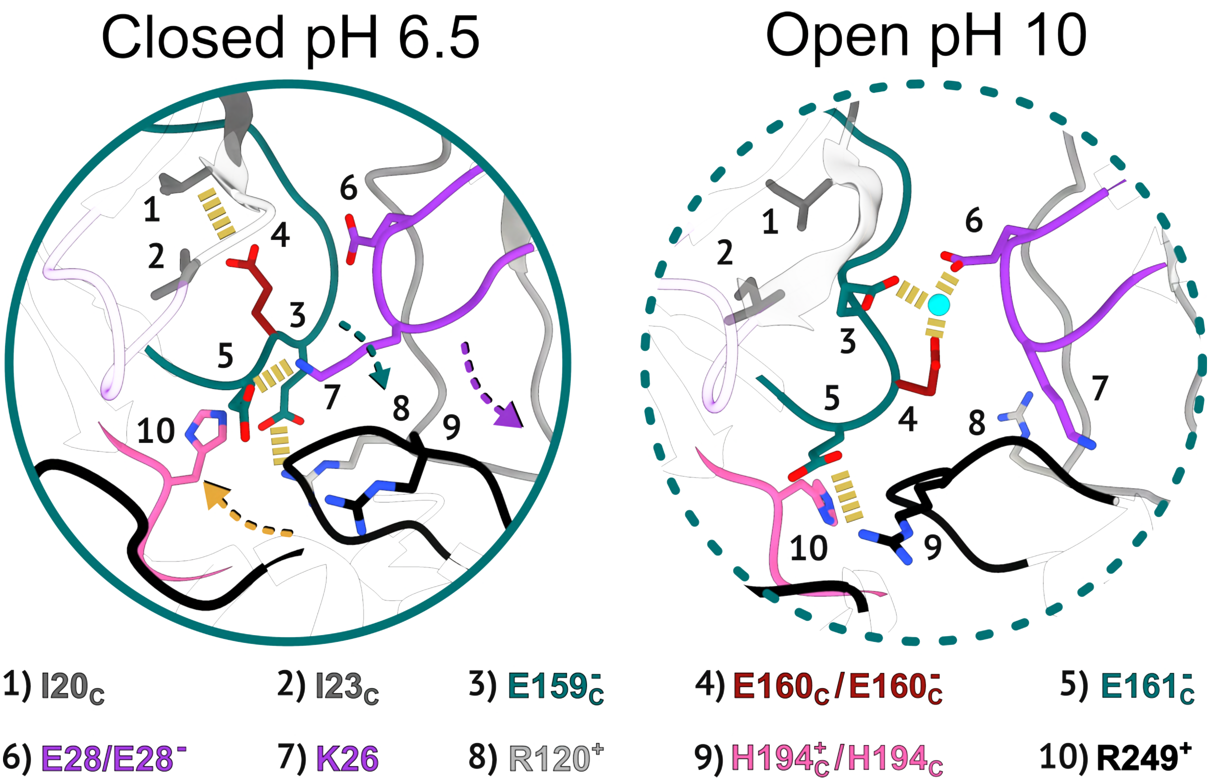


**Figure S4: Expanded representation of the ECD-TMD interface.** Zoom regions from simulations snapshots as in Fig. 3A, with additional structural details (numbered) in the closed state at pH 6.5 (left) and open state at pH 10 (right).

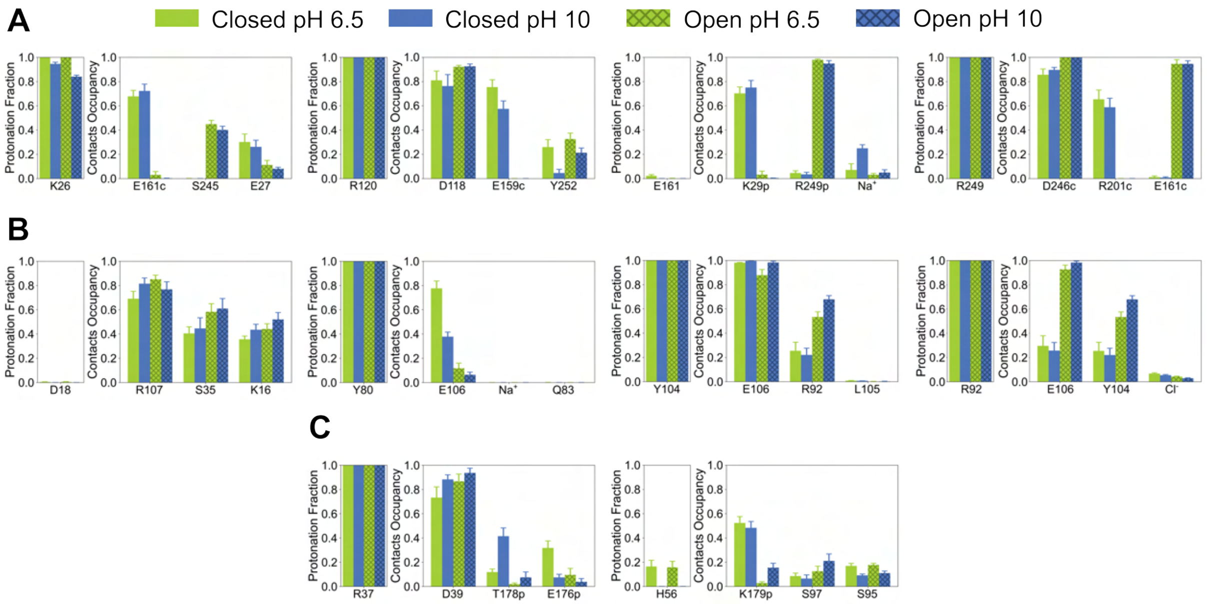


**Figure S5: Interaction details of additional residues potentially involved in pH gating. (A)** Mean protonation fraction and hydrogen-bond occupancies of (left to right) K26, R120, E161, and R249 at the subunit/domain interface. Columns represent simulations of the closed (solid) and open (crossed) states at pH 6.5 (lime) and pH 10 (blue), with error bars representing standard error of the mean. **(B)** Plots as in panel A for (left to right) D18, Y80, Y104, and R92 in the ECD vestibule. **(C)** Plots as in panel A for (left to right) R37 and H56 in the orthosteric site.

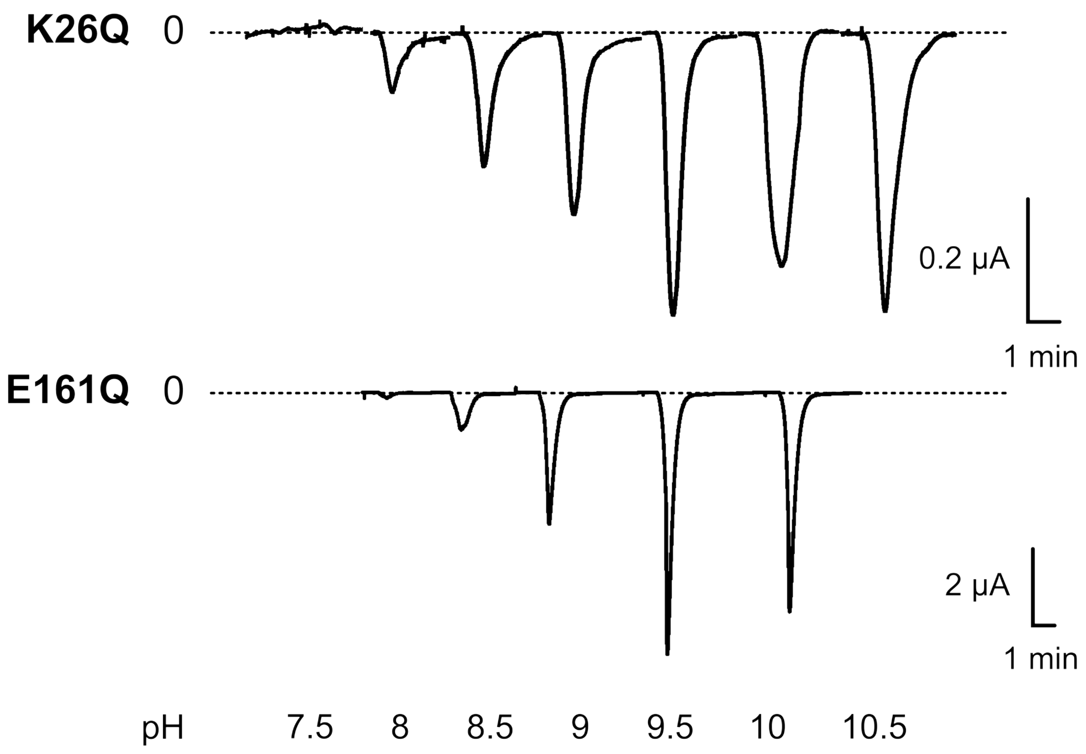


**Figure S6: Representative electrophysiology traces.** Current responses upon exposure to increasing pH for sTeLIC containing mutations K26Q and E161Q at the subunit/domain interface. The dashed line represents 0 μA current.

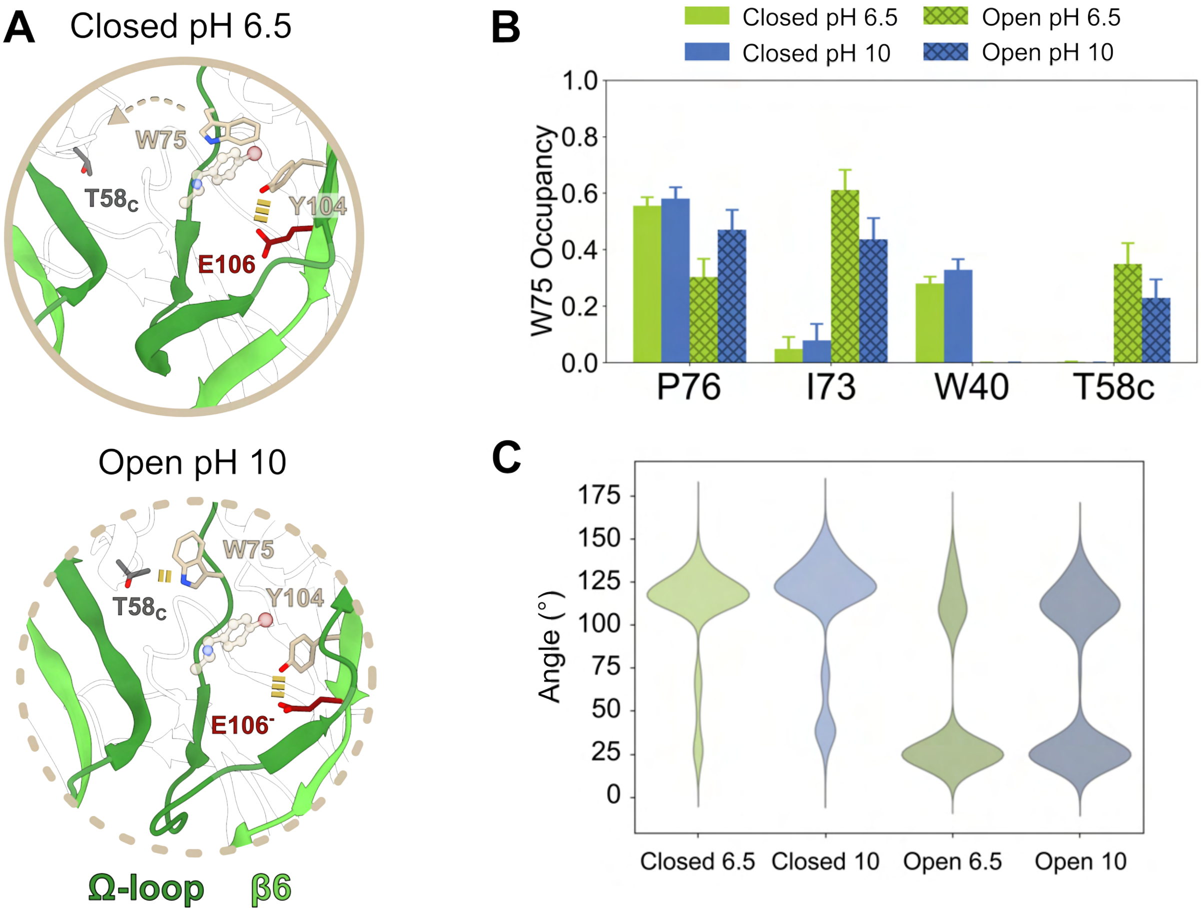


**Figure S7: State-dependent orientation dynamics of W75.** **(A)** Schematics as in Fig. 4A highlighting differential interactions of residues proximal to the vestibular modulation site [16] in the closed state at pH 6.5 (above, solid) and the open state at pH 10 (below, dotted). Key side chains, including E106 (red), are represented as sticks and labeled. For reference, 4-bromoamphetamine (balls and sticks) is superimposed from the open structure, wedged between W75 and Y104. Ribbons and side chains associated with the β4-β5 (Ω-loop) and β6 strands are colored dark and light green, respectively. Yellow dashes indicate state-specific polar contacts. **(B)** Hydrogen-bond occupancies of the most frequent contact partners for β4-W75. Columns represent simulations of the closed (solid) and open (crossed) states at pH 6.5 (lime) and pH 10 (blue), with error bars representing standard error of the mean. **(C)** Raincloud plot of W75 side-chain orientations across all subunits and simulations in a given condition, defined as the angle between two displacement vectors: one between the center of masses (COM) of two subunits, the other between the $C_\beta$ atom and imidazole-ring COM of W75.

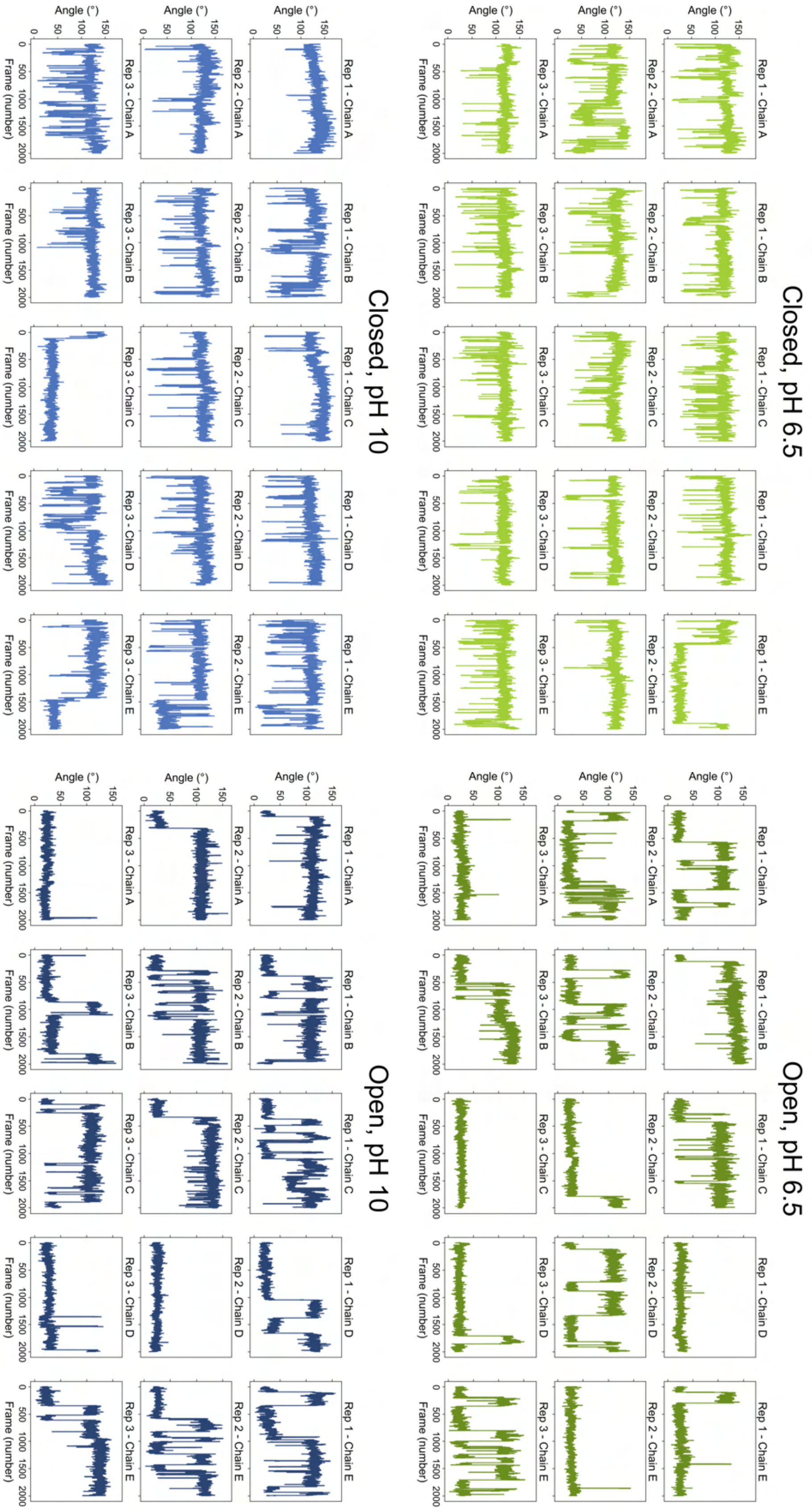


**Figure S8: Time-dependent orientation dynamics of W75.** W75 side-chain orientations for each condition, simulation (rows), and subunit (columns), defined as in Fig. S7C.

**Table S3: Partial charge parameters for deprotonated Tyr.** Charges were calculated using the CHARMM-GUI Ligand Modeler [36] (CE1, HE1, CZ, OH, HH, CE2, HE2). Parameterization of the correction potential followed the methodology described in [35], and the reference p$K_{\mathrm{a}}$ value was taken from [21].

| **CE1** | **HE1** | **CZ** | **OH** | **HH** | **CE2** | **HE2** |
|---|---|---|---|---|---|---|
| -0.600 | 0.280 | 0.400 | -0.760 | 0.000 | -0.600 | 0.280 |
